# Proteomic analysis reveals shared biological pathways linking acrolein to biomolecular changes in the acute phase of rat spinal cord injury

**DOI:** 10.64898/2026.03.11.711153

**Authors:** Rachel L. Stingel, Brendan K. Ball, Siyuan Sun, Douglas K. Brubaker, Riyi Shi

## Abstract

Spinal cord injury (SCI) pathology is highly difficult to treat due to substantial heterogeneity in injury presentation and spread, along with unclear mechanisms linking damage to pathology. Damages from injury forces (primary injury) are exacerbated by a series of biochemical events that follow the initial damage and injure additional tissue, known as secondary injury. Reactive aldehydes, such as acrolein, play a key role in propagating secondary injury cascades following SCI. Targeting acrolein after SCI has demonstrated therapeutic potential in limiting injury spread and pathology. However, injury mechanisms linking reactive aldehydes to SCI outcome have not been fully characterized. To gain a more comprehensive understanding of the cellular and molecular mechanisms underlying SCI, we generated proteomic profiles of rat spinal cords 24 h (acute phase) after subjection to SCI, sham injury, saline injection, or acrolein injection. We performed gene set enrichment analysis (GSEA) to characterize proteins and pathways significantly enriched after SCI and acrolein-injection. We then used Translatable Components Regression (TransComp-R), a framework for translating biological signatures across systems, to assess whether acrolein-associated spinal cord signatures can stratify SCI from sham outcomes. Our proteomics analysis revealed 467 differentially expressed proteins (DEPs) between the sham and SCI groups and 7 DEPs between saline and acrolein injection groups. Notably, the complement and coagulation cascades were upregulated in spinal cords subjected to SCI and acrolein injection. Our TransComp-R analysis further demonstrated that acrolein-associated signatures could distinguish SCI from sham conditions. Taken together, our findings suggest that acrolein induces proteomic alterations during the acute phase of SCI and is associated with complement and coagulation cascade activation, among other pathways. Therefore, this study reinforces the notion that understanding the role of acrolein in the acute phase of secondary SCI may be beneficial.

## INTRODUCTION

Spinal cord injury (SCI) is a devastating neurotraumatic event that often results in a profound and permanent loss of sensory, motor and autonomic functions. Although partial recovery is possible, outcomes are highly variable due to differences in injury severity and treatment factors.^1,2^ Furthermore, current therapies exhibit a limited ability to reverse the damage caused by SCI, thereby restricting rehabilitative potential.^3–6^ Improved understanding of SCI mechanisms will provide novel avenues to design more effective treatments.

Following the initial event causing SCI (primary injury), inflammatory and biochemical events (secondary injury), which can manifest in seconds and persist for years, exacerbate damage and worsen functional outcomes.^7^ While proteomics investigations have identified molecular mechanisms and pathways that are active during the secondary injury,^8–10^ the link between pathway alterations and injury pathology remains unclear. Furthermore, reliable biomarkers of secondary injury severity or progression remain elusive.

One group of promising molecular targets for SCI treatment and diagnosis are reactive aldehyde species such as acrolein.^11^ Starting in the acute phase of secondary injury (0-48 h), reactive aldehydes exacerbate oxidative stress, promote inflammation, induce structural damage, degrade myelin, and disrupt neuronal communication among other critical neurodegenerative pathologies.^11^ Acrolein, the most reactive α,β-unsaturated aldehyde, can bind to and modify DNA, proteins, and lipids causing structural changes and subsequent functional disruption.^12–16^ The formation of acrolein-protein adducts increases chemical reactivity and prolongs acrolein’s biological half-lives relative to free reactive aldehyde species.^12,13,15,17^ Consequently, acrolein can spread to sites distant from the primary injury.^12,15,18,19^ Spread and reactivity of acrolein has been shown to drive neuropathic pain^20–24^ and pathological protein aggregation associated with neurodegenerative pathologies^25–28^ all of which are long-term consequences following SCI. Despite these findings, the contributions of reactive aldehydes, such as acrolein, to proteomic changes in SCI progression and pathology have yet to be fully characterized.

This study investigated the link between acute secondary SCI and acrolein-associated pathology. Proteomics profiles were generated from rats given SCI, sham injury, acrolein injections, or saline injections. The resulting data were analyzed through a computational pipeline involving principal component analysis (PCA) and hierarchal clustering to identify proteins differentially expressed in each condition, gene set enrichment analysis (GSEA) to identify significantly enriched pathways, TransComp-R to assess the role of acrolein in driving expression changes after SCI, and Pathview to compare expression changes between shared genes that are key in acrolein and SCI pathways. In short, we identified shared proteomic responses to SCI and acrolein injection, which were associated with enriched pathways including the complement and coagulation cascades. Our use of TransComp-R further confirmed our findings that acrolein-associated signatures could indeed stratify SCI and sham outcomes, highlighting the importance of acrolein in SCI.

## MATERIALS AND METHODS

### Animal use and ethics statement

All animal studies were performed in strict accordance with the guidelines approved by the Purdue University Institutional Animal Care and Use Committee (IACUC) under protocol number 1111000095. This study is also reported in accordance with ARRIVE guidelines.

### Experimental design

Male Sprague Dawley rats were housed two per cage (200–300 g; Envigo RMS LLC) on a 12 h light-dark cycle and provided food and water *ad libitum*. All rats were acclimated to their surroundings for one week before surgical procedures and randomly assigned to one of the following groups: sham injury (laminectomy without SCI) or SCI for the first experiment or saline (vehicle control) or acrolein injection for the second experiment.

For surgical procedures, rats were anesthetized with an intraperitoneal mixture of ketamine (80 mg/kg) and xylazine (10 mg/kg) before undergoing a T10 laminectomy to expose the dorsal surface of the spinal cord. Rats in the first set of experiments were either subjected to a moderate spinal cord contusion SCI (force = 200 kDy) using a Horizon Impactor (Precision Systems and Instrumentation, Fairfax Station, VA) or received a laminectomy without SCI (sham). In the second set of experiments, the dura was removed following the laminectomy to allow for direct access to the spinal cord. A microinjector (model UMP3T-1, WPI, Sarasota, FL) mounted on a stereotaxic frame with a digital display (model 940, David Kopf Instruments, Tujunga, CA) was used to inject 1.6 µL of acrolein (40 nM, Oakwood Chemical) or saline (Aspen Veterinary Recourses) directly into the T10 spinal cord (coordinates: 0.6 mm from the midline and 1.3 mm from the dorsal surface, flow rate = 0.8 µL/min) (**Figure 1**).

**Figure 1.**
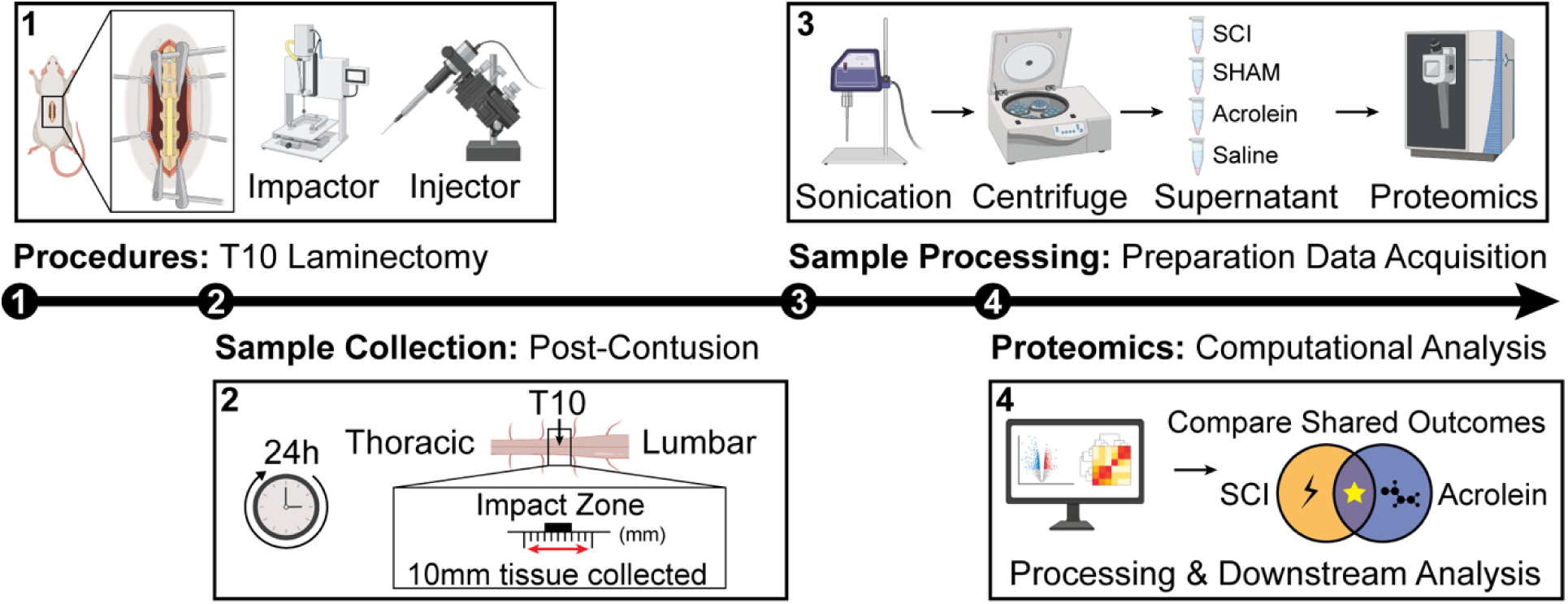
Experimental Overview. Rats were randomly assigned to sham, SCI, saline injection or acrolein injection groups. The SCI groups were impacted at T10 using the impactor while sham rats just received a laminectomy. Spinal cord samples were collected 24 h following procedures. Spinal cord samples were prepared, shipped to MtoZ Biolabs, and further prepared for liquid chromatography tandem mass spectrometry (LC-MS/MS). MtoZ Biolabs obtained readouts and returned the data for our downstream computational analysis. (Created with content from BioRender.com).

### Sample Preparation

Rats were sacrificed via transcardial perfusion with cold phosphate-buffered saline 24 h post-procedure. Next, 20 mg of spinal cord tissue containing the injury or injection site was excised and stored at -80 °C until shipment to MtoZ Biolabs (Boston, MA) for protein abundance quantification (**Figure 1**). At MtoZ Biolabs, spinal cord tissue, 200 µL of 8 M urea, and 2 µL of phenylmethylsulfonyl fluoride reagent were added to a grinding tube and homogenized. The supernatant was collected after centrifugation at 12,000 rpm for 10 min at 44 °C. For reduction and alkylation, 10 mM of dithiothreitol and 50 mM of ammonium bicarbonate were added to 50 µg of protein and incubated for 1 h at 56 °C. Next, 20 mM of Iodoacetamide was added and the solution was allowed to react in the dark at room temperature for 1 h. Lastly, the Iodoacetamide was neutralized by adding 10 mM of dithiothreitol and incubating for 1 h at 56 °C.

Beads (10μg/μL) were mixed with samples in a ratio of 10μg beads to 1μg of protein, vortexed, then incubated at room temperature for 10-15 min. The samples were then centrifuged and the supernatant was retained. The beads were cleaned 3 times with 80% ethanol then dried and resuspended in 50 mM of ammonium bicarbonate and shook in a metal bath at 1000 rpm for 5 min until they dispersed. Next, trypsin (0.25 μg/μL) was added, and the samples were allowed to digest overnight at 37 °C. The next morning, samples were placed on a magnetic rack and the supernatant was collected. The enzymatically hydrolyzed samples were then freeze dried using a lyophilizer. The samples were desalted using C18 desalting columns and dried with a vacuum centrifugal concentrator at 45 °C. Lastly, the samples were dissolved in 0.1% formic acid in preparation for mass spectrometry analysis.

### Liquid chromatography tandem mass spectrometry (LC-MS/MS)

The peptides were separated using Reprosil-Pur 120 C18-AQ 1.9 μm columns (150 μm i.d. × 170 mm). An elution gradient was created by mixing mobile phase solvent A (0.1% formic acid in aqueous solution) and solvent B (80% ACN, 0.1% formic acid). Solvent B was mixed into solvent A at gradually increasing concentrations: 5.8% for 1.8 min, 6.3% for 2 min, 23% for 56 min, 36% for 25 min, 55% for 1.5 min, and 99% for 7.5 min. The flow rate was set to 600 nL/min.

Mass spectrometry (MS) data was acquired using an Orbitrap Exploris 480 mass spectrometer (Thermo Fisher Scientific, Waltham, MA). The data-independent acquisition (DIA) full scan range was m/z 400-1100, with mass spectrometry stage 1 resolution at 120,000, automatic gain control at 300%, and maximum ion injection time at 22 ms. The mass spectrometry stage 2 was acquired with automatic gain control at 1000% and maximum injection time at 40 ms. Peptide fragmentation used collision energies of 25%, 27%, and 30%. The raw mass spectrometry data was then returned to us for analysis. In short, the mass spectrometer separates ions by mass-to-charge ratio (m/z) and quantifies peptide abundance (signal intensity) by measuring how many ions of each m/z (different m/z for each peptide/protein) hit the detector.

### Proteomics data processing and bioinformatics analysis

Peptides were identified from the raw mass spectrometry data with the Data-Independent Acquisition - Neural Networks (DIANN, v1.9) software using the UP000002494_10116.fasta. database. The following parameters were assigned for peptides: a maximum of two missed trypsin cleavages, 20 ppm peptide mass tolerance, and 20 ppm fragment mass tolerance. Fixed carbamidomethylation modifications and variable oxidation (M) and acetyl (N-terminus) modifications were also specified.

### Data pre-processing and normalization

All computational analysis was performed with R (v4.5.1). Data were split into two batches, based on experiment: (i) SCI vs. sham and (ii) acrolein vs saline injection. To reduce unintended bias from imputation on missing data, individual proteins missing more than 50% of entries prior to the downstream processing were removed. Each sample contained less than 2% missing data prior to data imputation (**Supplementary Table S1**). After pre-processing, the proteomics samples for each batch were log2 transformed and quantile normalized.

### Data imputation

It was first verified that all the samples contained missing data at random (**Supplementary Figure S1a,b**). Batches were imputed separately using the k-nearest neighbors (KNN) method (*k*=3) in R to estimate the missing entries based on similar neighboring samples (VIM v6.2.2).^29^ The KNN method identifies the *k* nearest samples using an Euclidean distance metric and imputes missing abundance values using the average values from its closest neighbors.^30^ After imputation, it was confirmed that the protein abundance distribution in both batches remained the same, indicating that the imputation method did not alter the data distribution (**Supplementary Figure S1c,d**).

### Differential protein abundance analysis

After imputing missing peptide abundances, differentially abundant proteins were identified using the R package limma (v3.58.1).^31^ A moderated t-test based on the empirical Bayes method was used to calculate p values to identify significant differences in peptide abundance. To account for multiple hypothesis testing, p values were adjusted using the Benjamini-Hochberg method, where a false discovery rate (FDR) less than 0.05 was identified to be statistically significant. Additionally, the limma package was used to calculate the log2 fold change (log2FC) in SCI in respect to the sham group or in the acrolein injection in respect to the saline injection group to determine directionality of changes in protein abundance. An absolute log2FC magnitude greater than 1 was set to identify proteins that had at least a two-fold difference in protein abundance between treatment (SCI or acrolein injection) and control (sham or saline injection).

### Principal component analysis and hierarchical clustering

The log2 transformed and z-score normalized proteomics data was used for principal component analysis (PCA) and hierarchical clustering across the SCI/sham and acrolein/saline injection batches (factoextra v1.0.7). Proteomics data are dimensionally reduced into a simplified set of variables called principal component (PC) vectors using PCA.

A volcano plot was employed to display significant changes in protein abundance between the treatment and control groups. Significant proteins with a Benjamini–Hochberg FDR of less than 0.05 and a log₂ fold change greater than 1 or less than negative 1 were filtered and labeled for hierarchical clustering. This approach allows for the identification of proteins with subtle changes that are statistically significant. A heatmap was used to visualize and compare samples and their respective protein abundances across conditions (pheatmap v1.0.12).^32^

### Gene set enrichment analysis

Identified peptides and log2FC values were employed in gene set enrichment analysis (GSEA) to identify differentially expressed pathways. To prepare for GSEA, UniProt identifiers were used to map gene expression symbols for *Rattus norvegicus*. The mapped genes were pre-ranked by their respective log2FC values using GSEA in R (fgsea v1.34.2 and clusterProfiler v4.16.0).^33^ Pathways had to have a minimum of 5 genes and could have a maximum of 500 genes. GSEA was run with the default of 1000 permutations, and the tuning constant, epsilon, was set to be 0 for the model parameters. Multiple data curations, Kyoto Encyclopedia of Genes and Genomes (KEGG), Hallmark, Reactome, and Gene Ontology: Biological Pathways) were accessed from the Molecular Signatures Database to identify enriched biological pathways (msigdbr v25.1.1). To account for multiple hypothesis testing, GSEA p values were adjusted into FDR using the Benjamini-Hochberg method.^34^ Significant pathways of interest associated under SCI and acrolein-injected conditions were identified as having an adjusted GSEA FDR value less than 0.05.

### Cross-condition modeling with TransComp-R

TransComp-R is a computational platform that allows for the comparison of transcriptomic data across species and conditions.^35^ Therefore, this method is ideal for assessing the contribution of acrolein to transcriptomic changes after SCI. Only peptides present in all four experimental groups (sham, SCI, acrolein injection, and saline injection) were included in TransComp-R analysis. TransComp-R was applied to the model by first creating a PCA with the acrolein/saline injection proteomic data. A dimensionally reduced matrix was created out of the PCs with a cumulative explained variance of 80%. This dimensionally reduced data is defined as matrix **Q**, with rows corresponding to the shared protein markers and columns as the acrolein-injected PCs. Next, the SCI vs sham rat proteomic data matrix **X** was projected onto the acrolein injection PC space, with rows corresponding to individual rat samples and columns to protein markers, by multiplying **X** × **Q**. The resulting matrix product, **P**, provides a synthesized matrix of the acrolein and SCI conditions, where the rows include the SCI and sham conditions, and the columns contain the acrolein-injected PCs.

### Selection of acrolein PCs predictive of SCI outcomes

Acrolein PCs predictive of SCI and sham conditions were identified by constructing a generalized linear regression model:

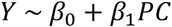

where **Y** represents the SCI or sham condition and **PC** is the acrolein vs. saline PC and its scores. A generalized linear model p value < 0.05 was established to determine acrolein PC’s ability to stratify SCI and sham outcomes.

### Principal component variance of acrolein conditions explained by SCI conditions

To determine acrolein’s contribution to SCI transcriptomic changes, the percent of variance in the SCI vs sham log2FC explained by acrolein vs. saline injection was quantified using the following equation:

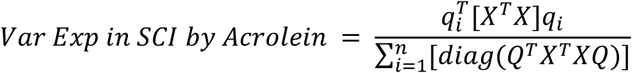

Where **X** is the SCI vs sham rat proteomic data matrix, **Q** contains the acrolein injection PCs, and **qi** is the ith PC in matrix **Q**. **T** represents the transpose of the matrix.

### Biomolecular pathway analysis visualization

Coagulation and complement cascades were consistently identified as enriched pathways in the GSEA and TransComp-R results. Pathview (v1.48.0)^36^ was used to compare mapped gene abundances driving signaling within the pathway. This approach allows the same pathway to be simultaneously compared under SCI and acrolein conditions.

To quantify the relationship strength between the SCI and acrolein injection conditions within the complement and coagulation cascade, the genes contributing to the pathway were identified with Pathview in R. A Pearson correlation analysis was performed using the log2FC of the identified proteins in both SCI and acrolein groups in respect to their sham and saline control conditions

## RESULTS

### Differentially expressed proteins between SCI and sham injuries

First, protein abundances between SCI and sham samples were analyzed to identify differentially expressed proteins. After pre-processing the SCI (n=5) and sham (n=5) data, the remaining 6,391 proteins were used for downstream computational analysis. To confirm that protein abundances varied between SCI and sham samples, a PCA was performed. The SCI and sham groups clustered separately from each other on PC1 with 39.3% of the variance explained (**Figure 2a**). Therefore, protein expression indeed varied between sham and SCI samples.

**Figure 2.**
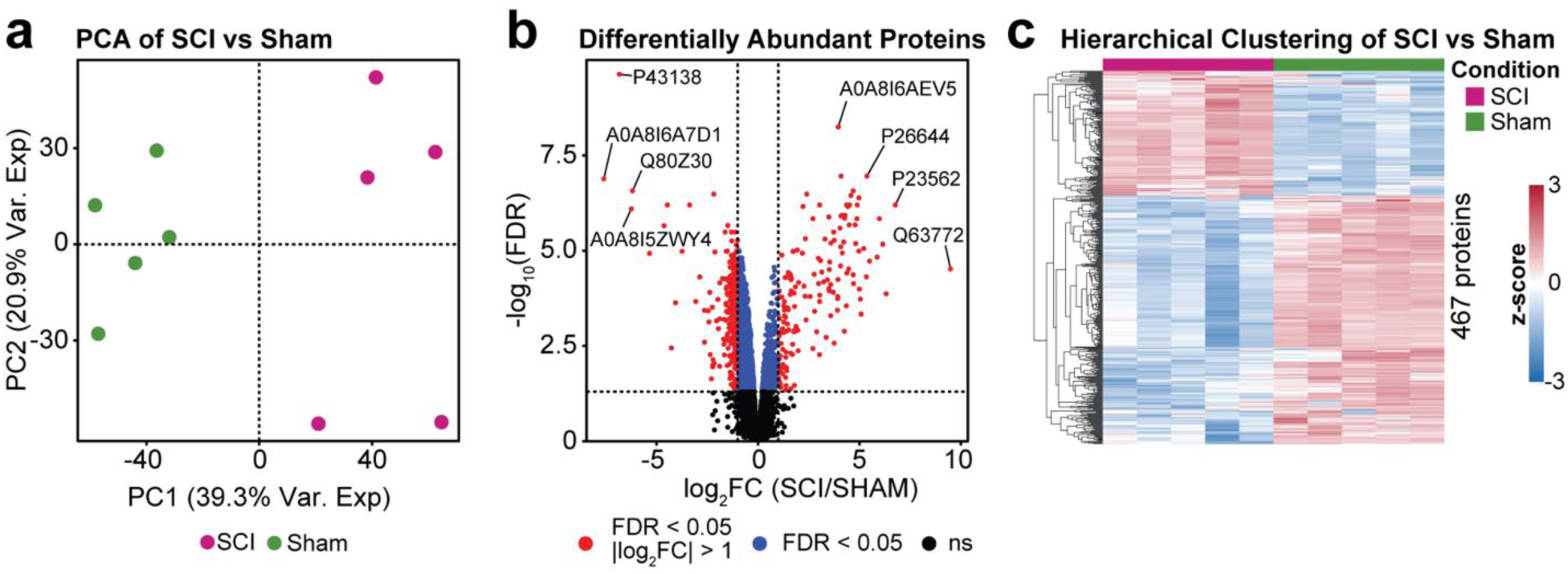
Differential abundant protein analysis comparing SCI and sham conditions. **(a)** PCA distinguishing the SCI and sham groups. **(b)** Differentially abundant proteins across SCI and sham groups, with the eight most extreme increases and decreases labeled by their UniprotID. **(c)** Hierarchical clustering analysis of the 497 proteins with an FDR value < 0.05 and SCI/sham |log2FC| > 1 was used to identify significantly upregulated and downregulated proteins.

Differential protein abundance analysis identified 2,567 proteins with an FDR less than 0.05. Of these, 467 also contained an absolute log2FC greater than 1 (**Figure 2b**). The four proteins most significantly increased after SCI were: Ig-like domain-containing protein (A0A8I6AEV5), beta-2-glycoprotein 1 (P26644), band 3 anion transport protein (P23562), and growth arrest-specific protein 6 (Q63772). Ig-like domain-containing protein is associated with immunoglobulins, which mediate the effector phase of the humoral immune response.^37^ Beta-2-glycoprotein binds to negatively charged entities (e.g., heparin, dextran sulfate), regulates the balance between the immune complement and coagulation systems, and may inhibit intrinsic blood coagulation by attaching to phospholipids on the surface of damaged cells.^38–40^ Band 3 anion transport protein facilitates CO2 transport and provides structural support in red blood cells.^41^ Growth arrest-specific protein 6 is actively involved in cell growth, survival, adhesion, migration, and inflammation.^42^ Together, these four most significantly upregulated proteins suggest that the greatest upregulation after SCI corresponds to promoting an immune response, altering blood clotting and red blood cell function, and modulating cell survival, adhesion, and migration.

The four most significantly downregulated proteins after SCI were: DNA repair nuclease/redox regulator APEX1 (P43138), ribosomal protein S6 kinase (A0A8I6A7D1), protein phosphatase 1E (Q80Z30), 2-aminoethanethiol dioxygenase (A0A8I5ZWY4). DNA repair nuclease/redox regulator APEX1 regulates transcription factors sensitive to oxidative stress and initiates DNA repair in response to oxidative stress.^43^ Additionally, APEX1 mediates Ig immunoglobulin class switch recombination.^44^ Ribosomal protein S6 kinase is involved in signal transduction of MAPK/ERK pathway.^45^ Protein phosphatase 1E is a protein that inactivates various substrates (i.e., PAK1, AMPK, CaMKs) via dephosphorylation.^46^ In humans, 2-aminoethanethiol dioxygenase regulates thiol metabolism and oxygen homeostasis.^47^ Together, these significantly downregulated proteins suggest that the greatest downregulation after SCI corresponds to reduced DNA repair in response to oxidative stress, reduced immune system Ig processing, and reduced or dysregulated kinase signaling.

Hierarchical clustering was performed with the goal of recognizing patterns and relationships in the data of the 467 differentially expressed proteins (DEPs). Of these proteins, 311 were downregulated and 156 were upregulated in SCI compared to sham (**Figure 2c**). Clear and coordinated similarities can been seen among samples with the same treatment whereas stark differences occur between groups. This observation reinforces the distinct clusters seen in the PCA and supports the notion that there are clear proteomic changes in response to SCI. All genes and their statistical outcomes are available in **Supplementary Table S1**.

### Differentially expressed proteins between acrolein and saline injections

Since acrolein is a key factor of secondary injury and potential therapeutic target, it was important to establish the contribution of acrolein alone in proteomic changes after SCI. To this end, it was necessary to identify differentially expressed proteins in spinal cords injected with acrolein compared to saline control. After proteomics pre-processing, 6,131 unique proteins were identified for downstream analysis across five saline and five acrolein-injected rats. The PCA found that the acrolein and saline injection samples clustered separately across both PC1 (49.5% variance explained) and PC2 (12.7% variance explained) (**Figure 3a**). Therefore, acrolein and saline injections elicited distinguished proteomic responses in the spinal cord.

**Figure 3.**
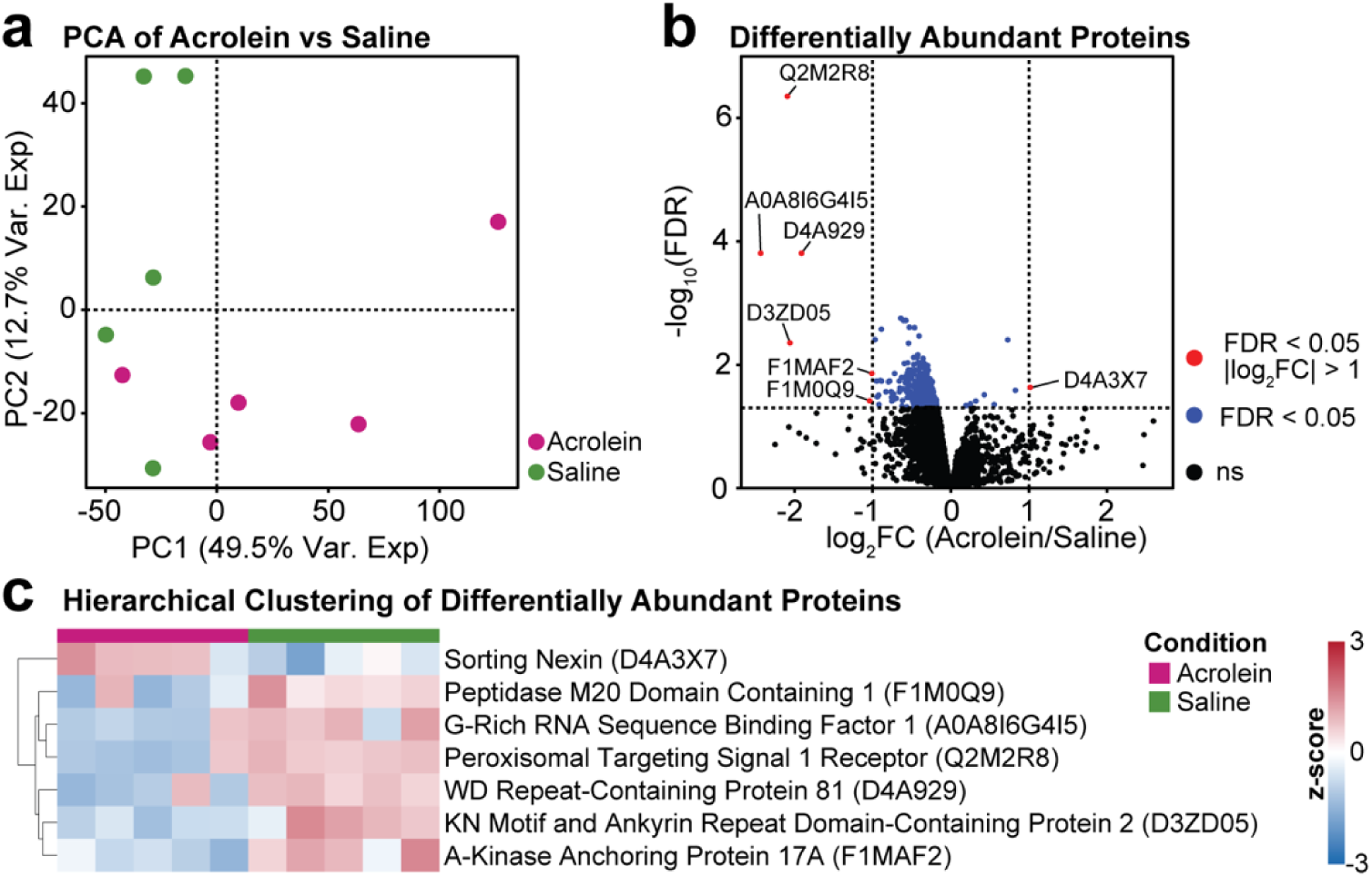
Differentially abundant proteins between acrolein and saline. **(a)** PCA distinguishing the acrolein and saline injected groups. **(b)** Differentially abundant proteins across acrolein and saline injected groups, with the seven most extreme increases or decreases labeled by their UniprotID. **(c)** Hierarchical clustering analysis of the most differentially abundant proteins under acrolein injected and saline, which contain 7 proteins with an adjusted FDR < 0.05 and acrolein/saline |log2FC| > 1.

Differential protein analysis identified 418 proteins with an FDR less than 0.05. Seven of these proteins also exhibited an absolute log2FC greater than 1 (**Figure 3b**). Only one of these seven proteins was upregulated. The upregulated protein, sorting nexin (D4A3X7), is a cytoplasmic protein that associates with membranes via its lipid-binding domain or by interacting with membrane proteins, and plays a role in protein trafficking.^48^ The other six proteins were downregulated and consist of: peptidase M20 domain containing 1, G-rich RNA sequence binding factor 1, peroxisomal targeting signal 1 receptor, WD repeat-containing protein 81, KN motif and ankyrin repeat domain-containing protein 2, and A-kinase anchoring protein 17A (**Figure 3b,c**). Peptidase M20 domain containing 1 (F1M0Q9) regulates amino acid metabolism and glucose homeostasis.^49^ G-rich RNA sequence binding factor 1 (A0A8I6G4I5) is important in mitochondrial function, RNA processing, and ribosome assembly.^50,51^ Peroxisomal targeting signal 1 receptor (Q2M2R8) is involved in peroxisomal uptake.^52^ WD repeat-containing protein 81 (D4A929) is thought to play a role in endo-lysosomal trafficking.^53^ KN motif and ankyrin repeat domain-containing protein 2 (D3ZD05) plays a role in regulating actin polymerization, cell signaling, and the cytoskeleton.^54^ A-kinase anchoring protein 17A (F1MAF2) is found in activated B-cells and regulates alternative splicing.^55^ Together, these proteins suggest that the greatest upregulation after acrolein exposure involves increased lipid binding interactions and altered protein trafficking. Conversely, the greatest downregulation after acrolein exposure corresponds to decreased metabolic function, decreased neuroprotection, decreased uptake in lysosome function, and decreased or altered signaling, gene expression, and cytoskeletal remodeling. All genes and their statistical outcomes are available in **Supplementary Table S2**.

### Proteins and pathways common to both acrolein and SCI are associated with immune, complement, and coagulation cascades

Of all proteins identified in the previous two sections, 144 contained an FDR < 0.05 in the SCI vs. sham and acrolein vs. saline batches. Of these 144 proteins, only one (G-rich RNA sequence binding factor 1, A0A8I6G4I5) was significantly altered (downregulated) by both SCI and acrolein (FDR < 0.05, |log2FC| > 1) (**Figure 2b, 3b, 4a**). G-rich RNA sequence binding factor 1 plays a role in post-transcriptional mitochondrial gene regulation as a component of the mitochondrial ribosome and is needed for recruiting mRNA and lncRNA.^51,56^ This common gene suggests that mitochondrial dysregulation contributes to commonalities between SCI and acrolein-induced injury.

Next, GSEA was used to identify biological pathways upregulated or downregulated in response to SCI and acrolein injection. The unfiltered gene list was pre-ranked by their log2FC values for GSEA. To identify consistently enriched biological pathways, four different data pathway curations were used: KEGG, Hallmark, Gene Ontology: Biological Process (GO:BP), and Reactome. The shared pathways that were significantly enriched (FDR < 0.05) in both the SCI and acrolein injection batches were compared. Under these four pathway analyses in GSEA, a positive normalized enrichment score (NES) indicates that a biological pathway is upregulated under a SCI or acrolein-injected conditions, whereas a negative NES suggests downregulation.

In the KEGG (**Figure 4b**) and Hallmark (**Figure 4c**) curations, the complement and coagulation cascades were identified as upregulated in both SCI and acrolein injection conditions. The complement and coagulation cascades regulate the innate immune response and clot formation to prevent excessive bleeding.

**Figure 4.**
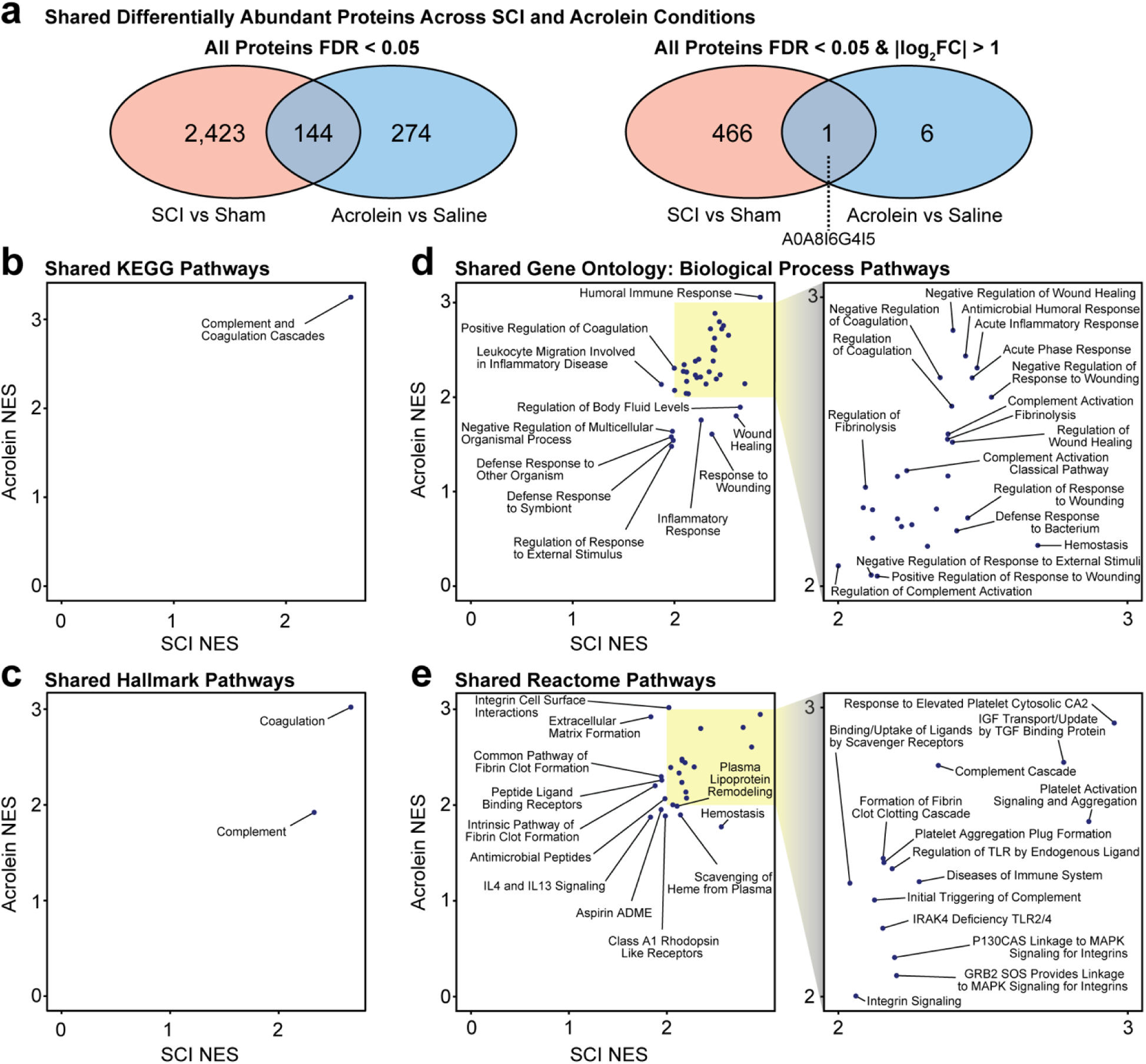
Shared enriched pathways across SCI and acrolein. **(a)** Common differentially abundant proteins across SCI and acrolein injection groups. **(b)** Shared GSEA KEGG pathways. **(c)** Shared hallmark pathways. **(d)** Shared GO:BP pathways. **(e)** Shared Reactome pathways. The yellow highlighted box indicates a zoomed-in cross-section of the plot. All pathways mentioned contain an FDR < 0.05 shared in both SCI and acrolein injection conditions.

From the biological processes database (GO:BP), 38 significantly enriched pathways were identified (**Figure 4d**). Notable pathways associated with both SCI and acrolein injection included: regulation of the complement and coagulation cascades, response to wounding, inflammation, and fibrinolysis. Likewise, 25 significantly enriched pathways were identified within the Reactome database (**Figure 4e**). These pathways regulate clotting, the complement cascade, and inflammatory signaling. All shared pathways identified by the four databases possess a positive NES score, indicating upregulation after SCI and acrolein exposure relative to sham or saline respectively. There were no downregulated pathways shared across SCI and acrolein injection groups in any of the four GSEA curations. All pathways prior to cross-condition matching identified by GSEA are also available in the supplementary material: KEGG (**Supplementary Tables S3-4**), Hallmark (**Supplementary Tables S5-6**), GO:BP (**Supplementary Tables S7-8**), and Reactome (**Supplementary Tables S9-10**).

### Translational modeling identifies gene-mapped expression changes due to acrolein injection predictive of SCI vs. sham

After separately analyzing the SCI and acrolein injection data, we were determined to test if the proteomic changes induced by acrolein injection can stratify the protein abundance changes under SCI and sham conditions. TransComp-R was leveraged to identify transcriptomic changes due to acrolein that might drive changes after SCI. TransComp-R has been successfully employed to identify causative transcriptomic changes and potential therapeutic targets in the context of colon cancer and colitis,^35,57^ Alzheimer’s disease,^58^ osteoarthritis^59^ and space-induced muscle loss.^60^

The TransComp-R pipeline works by first matching for shared proteins which are mapped into gene symbols (**Figure 5a**). Next, a PCA space is created containing the variance among the acrolein and saline injection samples before the SCI and sham samples are projected into this dimensionally reduced space. The combined data were regressed against SCI and sham outcomes, and acrolein principal components that significantly distinguished between the SCI and sham conditions were selected. The biological signal encoded on the significant PCs were interpreted using computational analyses such as GSEA.

**Figure 5.**
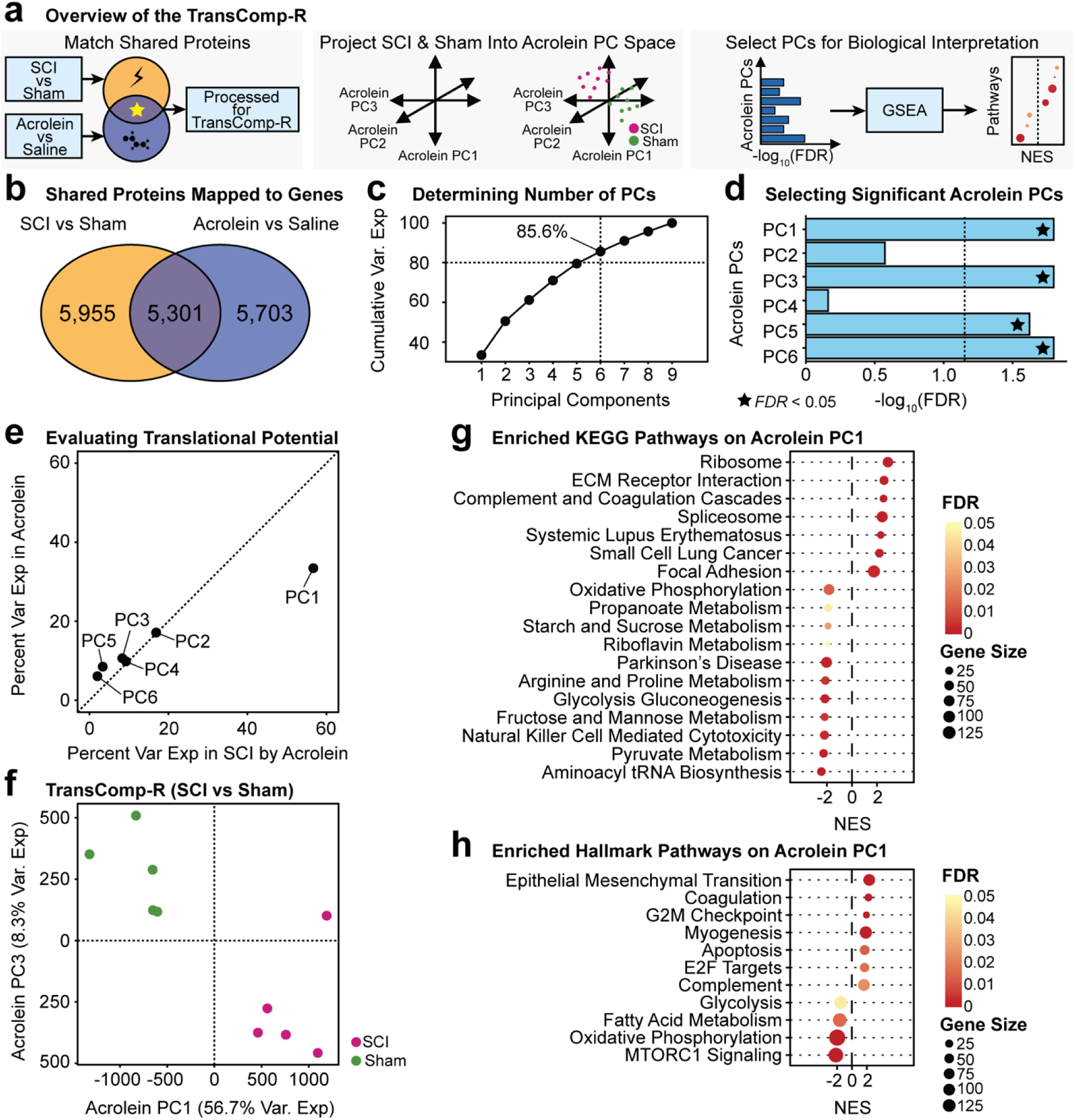
TransComp-R identification of acrolein gene mapped changes predictive of SCI. **(a)** Overview of the TransComp-R methodology. The log2FC of proteins mapped to gene symbols and common to both SCI and acrolein injection batches are used to project SCI vs. sham into acrolein vs. saline. Next, a matrix of acrolein vs. saline PCs is constructed before a matrix of SCI and sham samples are projected into the acrolein PC space. Linear regression is then used to identify acrolein PCs that can stratify SCI vs sham conditions. These acrolein PCs are then selected for GSEA to identify relevant biological pathways. **(b)** A total of 5,301 shared proteins were mapped to genes across the two data groups. **(c)** Cumulative variance explained to determine the number of relevant acrolein vs saline PCs. **(d)** Selection of significant acrolein PCs is predictive of SCI vs. sham determined by linear regression. Significance determined by a Benjamini-Hochberg adjusted FDR < 0.05. **(e)** Variance explained by the acrolein PCs for both acrolein (from PCA) and SCI. **(f)** Principal component plots of two acrolein PCs that both separate SCI and sham conditions (panel d) and explain a large percentage of the variance in both batches (panel e). **(g)** KEGG GSEA pre-ranked by the scores of acrolein PC1. **(h)** Hallmark GSEA pre-ranked by the scores on acrolein PC1. A positive NES in both KEGG and Hallmark indicates SCI signatures predicted by acrolein.

Of the proteins mapped to gene symbols, 5,301 genes were shared across the rat conditions for translational modeling (**Figure 5b**). For the TransComp-R model, six acrolein PCs were identified that explained at least 80% of the cumulative variance (**Figure 5c**). Linear regression on the acrolein PCs projected with the SCI/sham groups identified PC1, PC3, PC5, and PC6 as significant (p value < 0.05) in predicting SCI vs. sham.

The extent to which the acrolein PCs explained the SCI data was calculated to determine which significant PCs accounted for the greatest variance in the SCI and sham batches. Of the six PCs, PC1 had the most translational potential into the SCI data due to the higher proportion of variance explained in SCI by acrolein compared to the other PCs (**Figure 5d**). Likewise, distinct separation among the SCI and sham conditions on acrolein PC1 was seen (**Figure 5e**). Because of this finding, acrolein PC1 was used for downstream biological pathway analysis.

GSEA identified 18 enriched KEGG (**Figure 5g**) and 11 Hallmark (**Figure 5h**) acrolein injection pathways predictive of SCI. Notably, the complement and coagulation cascades were upregulated, and oxidative phosphorylation was found to be downregulated (**Figure 5f**). Oxidative phosphorylation is the final step in cellular respiration and the step where ATP is generated. Another notable downregulated pathway from the Hallmark GSEA is mTOR signaling; dysfunction of this pathway has been linked to neurodegeneration.^61^

### Similar complement and coagulation cascade activation after SCI and acrolein injection

Both GSEA and TransComp-R indicated upregulation of the complement and coagulation cascades after SCI and acrolein injection. Importantly, acrolein-induced inflammation and inflammation-induced acrolein production form a pathological cycle that can play a key role in driving secondary injury propagation. To further investigate the link between acrolein, injury, and inflammation, Pathview^36^ was employed to study more detailed, gene-level alterations in the complement and coagulation cascade. Pathview allows for direct comparison of genes and pathway interactions after SCI (**Figure 6a**) and acrolein injection (**Figure 6b**). Gene expression changes in both conditions shared similar directionalities, and most of the affected genes were elevated after SCI or acrolein injection. Genes that shared similar directionality in fold change included A2M, C1qrs, C3, C4, cofactor, FH, fibrinogen, KNG, MAC, PLG, and serpin.

**Figure 6.**
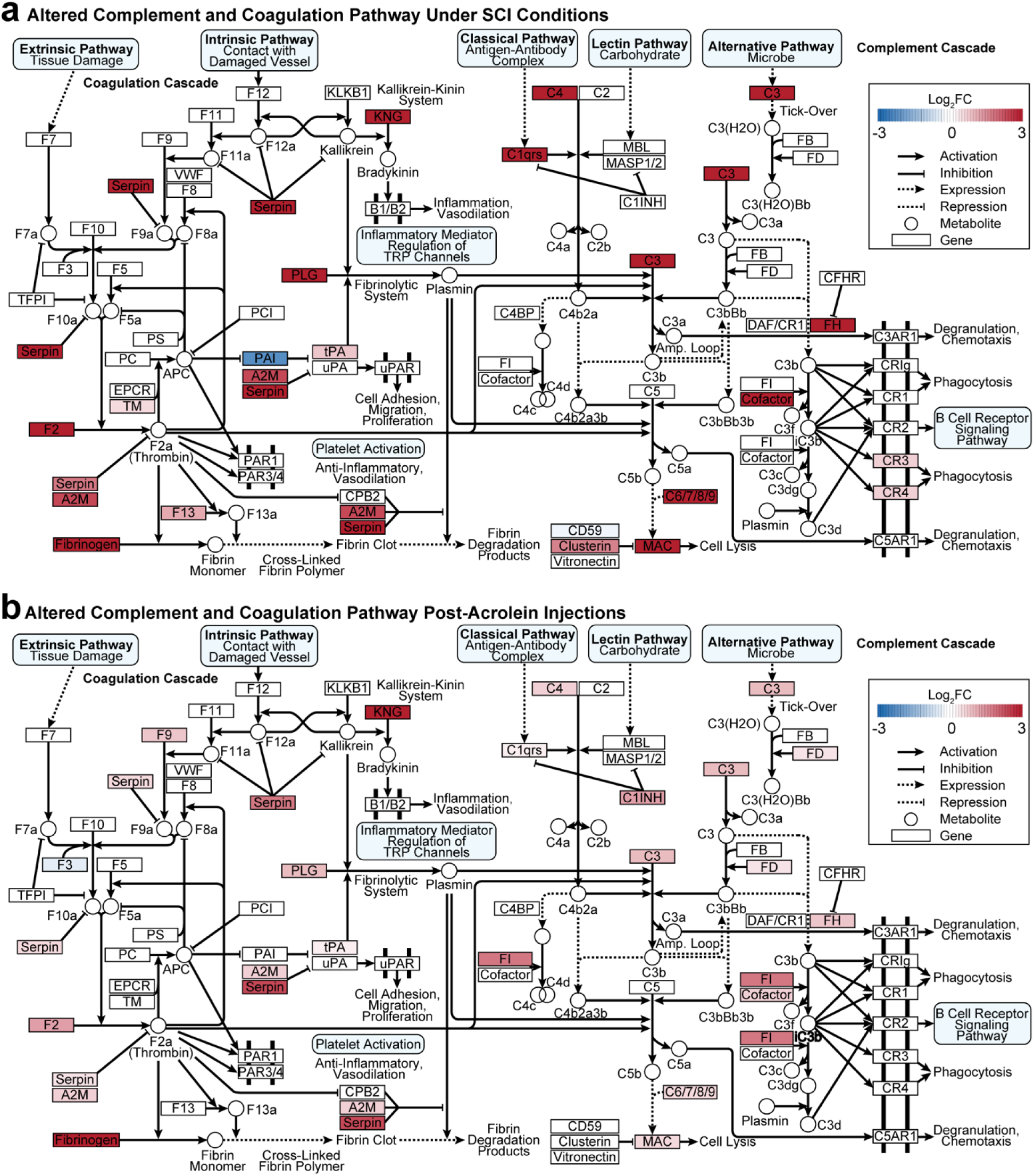
Mapped alterations of the complement and coagulation pathway due to acrolein or SCI. **(a)** Altered pathway genes under SCI conditions. **(b)** Altered pathway genes post-acrolein injection.

The results in **Figure 6** suggest that the same genes are affected in a similar manner in SCI and acrolein injection. A Pearson correlational analysis was performed to confirm this finding quantitatively. Using Pathview, 10 shared genes were identified between the SCI and acrolein injection that contributed to pathway enrichment (**Figure 7a**). These genes included Cd59b, Cfh, Clu, F13a1, F2, F3, Plat, Plg, Serpine2, and Serping1. Of the 10 genes, F2, Plg, Serping1, Cfh, Plat, and Clu were elevated under SCI or acrolein exposure, whereas Serpine2, Cd59b, and F3 abundance were reduced.

**Figure 7.**
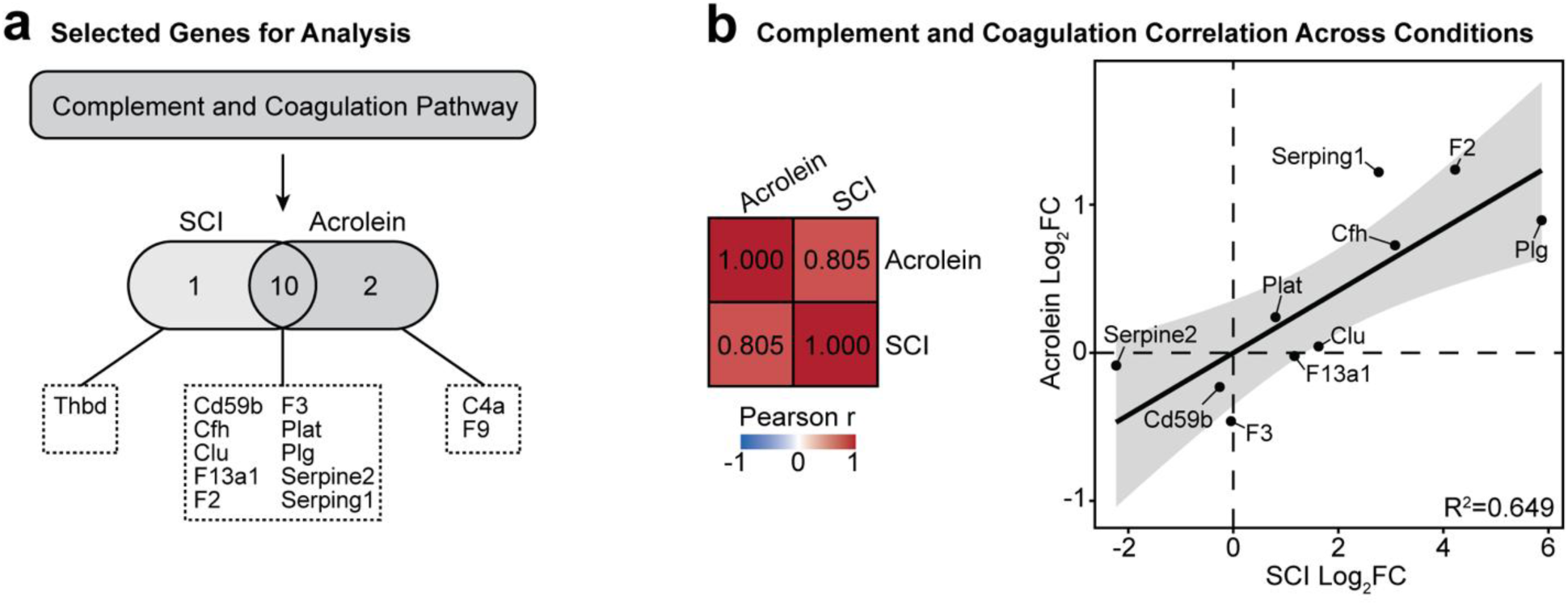
Pathway similarity analysis across SCI and acrolein injection proteomic abundances. **(a)** Selection of shared protein-coding genes from the complement and coagulation pathway used for analysis. **(b)** Pearson correlation analysis and linear regression model of the genes and their respective log2FC responses in acrolein against SCI. The linear fit is represented by the shaded standard error range, with R^2^=0.649.

Another Pearson correlation analysis was performed, this time between the log2FC of the acrolein injection and SCI batches relative to their respective control groups. It was determined that the 10 genes and their abundances contained a moderate correlation of 0.805 between the two conditions (red matrix, **Figure 7b**). This relationship was further visualized with a direct relationship between each of the expression biomarkers (R^2^=0.649, **Figure 7b**). The moderate correlation suggests that acrolein may interact directly or indirectly with these proteins after SCI, leading to alterations in gene expression. This change in gene expression could have an impact on downstream secondary injury pathology.

## DISCUSSION

Acrolein is a highly reactive aldehyde involved in the secondary injury after SCI. However, there is an incomplete understanding of changes on the proteomics level that can be attributed to acrolein, especially in the acute phase of injury. In this work, we leveraged computational approaches, including unsupervised hierarchical clustering, GSEA, TransComp-R, and Pathview, to characterize the proteomic profiles of spinal cords in the acute phase (24 h) following SCI or acrolein injection. Our analysis revealed consistent upregulation common between SCI and acrolein involved the complement and coagulation cascade. This commonality was observed from the protein level (PCA), the pathway-level (GSEA), and the shared pathway-level (TransComp-R). Furthermore, alterations to the complement and coagulation cascade were moderately correlated between SCI vs. sham, and acrolein vs. saline, suggesting that acrolein modulates these pathways through a combination of direct and indirect effects after SCI.

Differential expression analysis along with hierarchal clustering methods were employed to determine that 497 proteins were differentially abundant in SCI vs. sham, with 156 upregulated and 311 downregulated. Clustering of these 497 proteins distinguished SCI vs sham. Similarly, 7 proteins were differentially abundant in acrolein vs saline, with 1 upregulated and 6 downregulated. The fewer differentially abundant proteins for acrolein vs. saline compared to SCI vs. sham may arise due to a few reasons. First, acrolein-protein adducts peak around 24 h post-SCI but levels can remain elevated for more than 4 weeks following injury.^62^ The 24 h post-injury/injection may have been too early for acrolein-induced effects to fully accumulate, as reported in other papers from our group.^19,21,24,63^ However, this is unlikely because GSEA, TransComp-R, and Pathview analysis demonstrated that acrolein transcriptomic changes already relate to those observed after SCI. Therefore, timeline effects, while possible, do not completely explain the difference in significantly altered abundances between the treatment batches. A second explanation is that acrolein-induced changes, while critical, are likely only a subset of the changes induced by SCI and therefore would exhibit less of an effect than all SCI factors combined. For example, secondary injury can also be spread through pro-inflammatory signaling, damaged blood vessels, and excitotoxicity.^3,64^

Of the proteins we analyzed, we found one (G-rich sequence factor 1, A0A8I6G4I5) that was differentially expressed and downregulated in both the SCI and acrolein conditions. Another study also found that G-rich sequence factor 1 expression decreases following SCI. ^65^ The same study also found that overexpression of G-rich sequence factor 1 attenuates ferroptosis,^65^ cell death due to the accumulation of lipid peroxides. Therefore, G-rich sequence factor 1 can be neuroprotective and downregulation may render cells more susceptible to neurodegenerative changes. Furthermore, since G-rich sequence factor 1 protects against ferroptosis, reduction in this protein after SCI and acrolein injection might render cells more sensitive to acrolein, which promotes and is generated by lipid peroxidation.^66^

Several of the differentially expressed proteins in SCI vs. sham or acrolein vs. saline, while not common between the batches, operated downstream or in parallel with the complement and coagulation cascades. Glutathione peroxidase 1 acts upstream by reducing oxidative stress,^67^ which directly and indirectly diminishes coagulation and complement activation.^68^ Likewise, glutathione S-transferase theta-1 helps maintain redox balance,^69^ thereby indirectly limiting oxidative conditions known to amplify both complement activation and pro-coagulant signaling. Proteins involved in platelet-activating factor metabolism, such as platelet-activating factor acetylhydrolase IB subunit α1, act upstream as a potent inducer of platelet activation, vascular permeability, and inflammatory responses^70^ that potentiate hemostasis and thrombosis. Tight junction protein ZO-1 influences endothelial barrier integrity^71,72^ and thus tissue-factor exposure, contributing indirectly to the regulation of environments in which complement and coagulation operate.

While differences in protein abundance may alter immune activity, oxidative stress, cell gene expression, migration, and metabolic changes, it is necessary to validate these notions. Additionally, it is possible that significant differences in protein abundances are canceled by other, more subtle protein expression changes in the affected signaling pathways. Therefore, the full effects of changes in individual protein abundances, as well as how acrolein might relate to SCI, can only be analyzed by assessing the impact on overall signaling pathways and physiological processes. GSEA was employed to perform this assessment.

Despite fewer significant changes in protein abundance, acrolein injection significantly altered a variety of pathways. In both SCI and acrolein injection conditions, the complement and coagulation cascades were consistently identified to be active and upregulated, which is corroborated by other independent studies.^8^ Complement activation occurs at the injury site within 24 h of neurotrauma.^73,74^ This activation is facilitated by disruption of the blood-spinal cord barrier, which permits the leakage of circulating blood components into the central nervous system.^75^ Complement proteins may enter through the compromised blood-spinal cord barrier or be produced locally and released by infiltrating immune cells and resident cells.^73^ Once present, complement and coagulation cascade signaling can further exacerbate barrier dysfunction, promoting increased permeability and amplifying the influx of inflammatory cells and additional complement factors.^76^ Complement signaling can drive downstream pathological processes, including immune cell recruitment, myelin clearance, cytokine production, and cell death.^73^ Accordingly, dysregulation of the complement and coagulation cascades is well recognized as a detrimental pathology and has been implicated in both traumatic injury and disease states.^76^

While GSEA identified that the complement and coagulation cascades were upregulated in both SCI vs. sham and acrolein vs. saline, it remained unclear whether acrolein might drive this regulation after SCI. TransComp-R was used to overcome this limitation and assess whether acrolein might drive coagulation and complement cascade upregulation, or exert effects on additional pathways after SCI.

A notable finding from our TransComp-R model showed that proteomic variance across acrolein and saline injections could predict sham vs. SCI. This further validates that acrolein serves an active role in driving neurodegenerative pathology in the acute phase of secondary SCI. This is in combination with previous studies that have demonstrated acrolein scavenging increased neuroprotection and functional recovery following SCI. ^62,77,78^ Reactive aldehydes like acrolein arise after SCI through oxidative stress and lipid peroxidation, and further amplify these same processes in a pathological feed-forward cycle.^66^ Acrolein is capable of redox dysregulation, direct tissue destruction, inducing pro-inflammatory signaling, damaging mitochondria, and directly binding to/damaging DNA and proteins.^14,79–82^ Downstream, these processes lead to neurodegeneration, neuropathic pain and neuronal cell death.^83^ Acrolein has also been implicated in neurodegenerative diseases such as Parkinson’s Disease^25^ and Alzheimer’s Disease.^28^ In addition to these previously characterized pathologies, our data also implicates acrolein in modulating the coagulation and complement cascades. Ultimately, acrolein possesses a clear and highly destructive role in neurodegenerative pathology, which motivated us to characterize its role in proteomic changes following SCI. While acrolein has already been shown to promote inflammation after SCI,^84,85^ our results implicate this promotion in driving transcriptomic changes after SCI.

GSEA performed on acrolein PC1 identified by TransComp-R, as explaining the highest amount ∼60%) of the variance in SCI vs. sham log2FC, implicated acrolein in a variety of signaling effects after SCI. After SCI, pathways upregulated by acrolein involved altered gene translation (ribosome, spliceosome), progression through the cell cycle (G2M checkpoint, E2F targets), altered cell migration or function (ECM receptor interaction, focal adhesion), cell phenotype changes that promote wound healing (epithelial mesenchymal transition), and pro-inflammatory responses (complement and coagulation cascades). Additionally, pathways identified such as ribosome and cell migration also relate to the immune response. For example, ribosome function and splicing plays a key role in processing antibodies that dictate the immune response. ECM receptor interactions and focal adhesions play a key role in immune cell trafficking. Together, these results implicate a variety of ways acrolein might enhance the complement and coagulation cascades.

After SCI, acrolein appears to be involved in the downregulation of pathways related to metabolism. Reduced metabolism has been implicated in neurodegenerative pathologies.^86^ Other pathways suggest dampened efficacy of the innate immune response, suggesting that transcriptomic damage from acrolein may not only promote inflammation, may also render immune cells less effective at target elimination. This further implicates the role of acrolein in inflammatory dysfunction following SCI.

To further explore the pathological cycle in which acrolein promotes inflammation and inflammation promotes acrolein, Pathview was used to study how individual genes in the complement and coagulation cascade were altered by both SCI and acrolein injection. We found considerable similarities in the directionality of protein expression in both conditions, which implicates acrolein pathology in these changes following SCI. Genes that shared similar directionality in fold change included A2M (Alpha-2-Macroglobulin), C1qrs (Complement Component 1q), C3, C4, FH (Factor H), fibrinogen, KNG (Kininogen-1), MAC (Membrane Attack Complex), PLG (plasminogen), and serpin. Serpin family B member 9 (Serpin B9,Q6AYF8) was found to have an FDR less than 0.05 in both the SCI and acrolein injected conditions and a negative log2FC. Serpin B9’s chief function is protecting immune cells from programmed apoptosis through granzymeB.^87^

This work raises several future directions for understanding how acrolein drives SCI pathology. First, only male rats were used in our study because males represent 78% of SCI cases.^88^ Further studies on the female population in SCI may uncover additional or different mechanisms by which acrolein drives SCI pathology. Secondly, the TransComp-R model is only capable of selecting shared proteins across both treatment batches; therefore, there is a possibility that some biomarkers may have been omitted from TransComp-R analysis. Similarly, a minor portion (6.8% for SCI and 7.0% for acrolein) of the total identified proteins did not map to genes for GSEA, thus some pathways may have slight variances in significance. Third, the methods used for sample processing, although well established and highly utilized in proteomics, are not optimized to capture membrane proteins. Membrane proteins are notoriously difficult to process and obtain results due to their partially hydrophobic surfaces and difficulty to digest.^89,90^ Thus, while most of the proteome was mapped in this study, little if any transmembrane proteins were included in the data set. In spite of these limitations, this study still improves our understanding on the mechanisms by which acrolein promotes proteomic and transcriptomic changes after SCI.

This study highlights potentially altered pathways after SCI and a simulated secondary injury with acrolein. Our systems biology analysis suggested that acrolein may play a key role in upregulating the complement and coagulation cascade in the acute phase of injury. In addition to these changes, changes to specific genes indicate additional links between acrolein and secondary injury sequelae. These findings further improve our understanding of the acute effects and biological changes under a SCI and can therefore be used to investigate new therapeutic avenues for SCI and other diseases where acrolein has been shown to play a critical role.

## ACKNOWLEDGEMENTS

RLS is supported by the Bilsland Dissertation Fellowship. BKB acknowledges the National Science Foundation for support under the Graduate Research Fellowship Program (DGE-1842166). RS is supported by the National Institutes of Health, grant number R21NS115094 and the State of Indiana. The authors additionally thank Casey Adam for manuscript proofreading and editing. MtoZ biolabs was responsible for proteomic sample processing.

## AUTHOR CONTRIBUTIONS

**RLS:** Conceptualization, data curation, formal analysis, investigation, methodology, visualization, funding acquisition, writing-original draft, writing-review & editing. **BKB:** Conceptualization, data curation, formal analysis, investigation, methodology, visualization, funding acquisition, writing-original draft, writing-review & editing. **SS:** Data curation, methodology, writing-review & editing. **DKB:** Project administration, resources, writing-review & editing. **RS:** Conceptualization, funding acquisition, project administration, resources, writing-review & editing.

## DATA AND CODE AVAILABILITY

All proteomics data and code used from the study is deposited and made publicly available on https://github.com/bbkazu5/SCI-Proteomics.

## SUPPLEMENTARY FIGURES

**Supplementary Figure S1.**
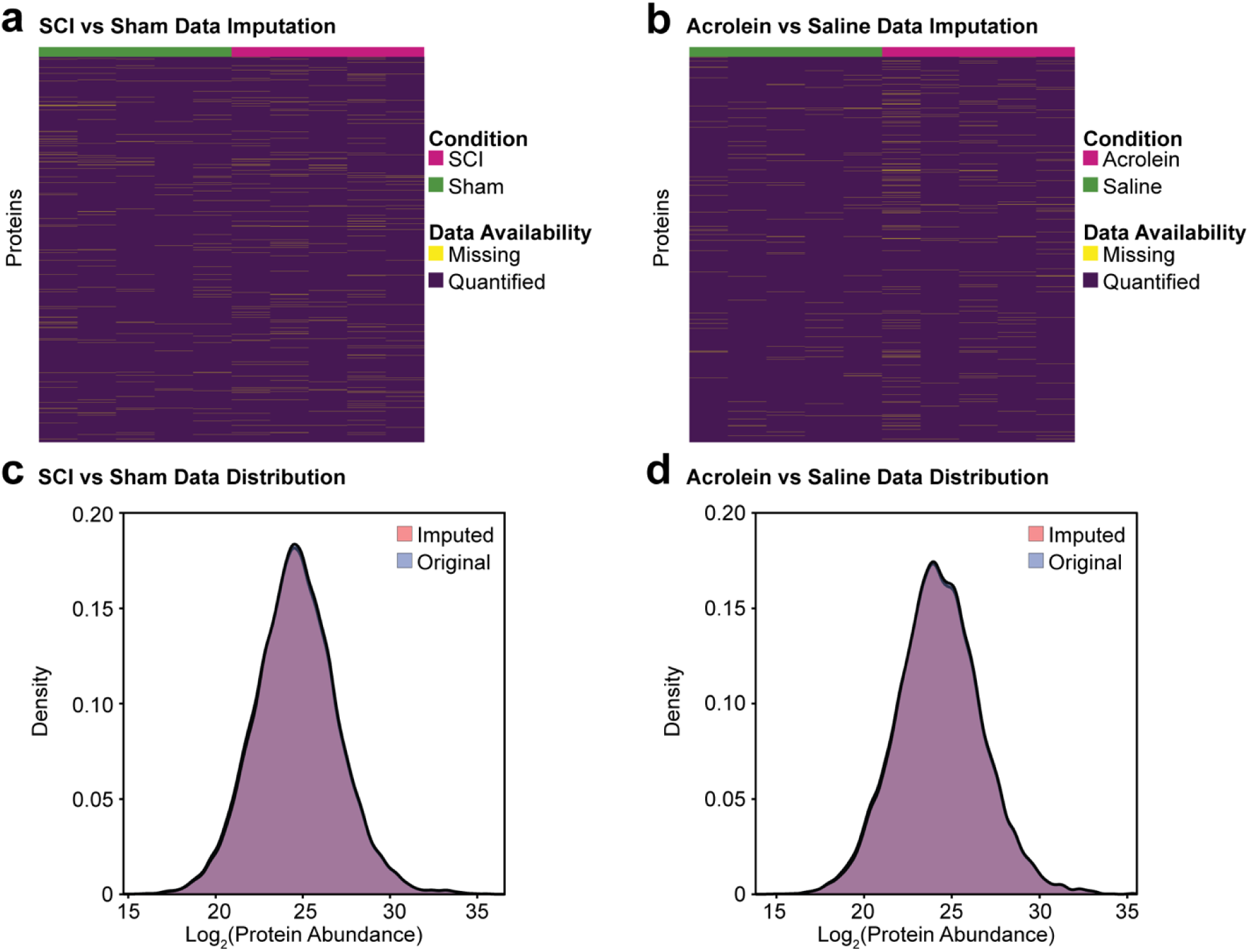
Pre-processing of the proteomics data. **(a)** Data imputation of SCI and sham data. **(b)** Data imputation of the acrolein and saline injection data. **(c)** Data distribution pre- and post-imputation for SCI. **(d)** Data distribution pre- and post-imputation for acrolein.

